# Silent transcription intervals and translational bursting lead to diverse phenotypic switching

**DOI:** 10.1101/2022.08.04.502777

**Authors:** Xiyan Yang, Songhao Luo, Zhenquan Zhang, Zihao Wang, Tianshou Zhou, Jiajun Zhang

**Author notes:** Author to whom correspondence should be addressed. Electronic address.

## Abstract

Bimodality of gene expression, as a mechanism generating phenotypic diversity, enhances the survival of cells in a fluctuating environment. Growing experimental evidence suggests that silent transcription intervals and translational bursting play important roles in regulating phenotypic switching. Characterizing these kinetics is challenging. Here, we develop an interpretable and tractable model, the generalized telegraph model (GTM), which considers silent transcription intervals described by a general waiting-time distribution and translational bursting characterized by an arbitrary distribution. Using methods of queuing theory, we derive analytical expressions of all moment statistics and distribution of protein counts. We show that non-exponential inactive times and translational bursting can lead to two nonzero bimodalities that cannot be captured in the classical telegraph model (CTM). In addition, we find that both silent-intervals noise and translational burst-size noise can amplify gene expression noise, as well as induce diverse dynamic expressions. Our results not only provide an alternative mechanism for phenotypic switching driven by silent transcription intervals and translational bursting, but also can be used to infer complex promoter structures based on experimental data.

**SIGNIFICANCE:** Understanding how phenotypic diversity arises among isogenic cell populations is a fundamental problem in biology. Previous studies have shown that the bimodality of gene expression contributing to phenotypic variations is mostly caused by the intrinsic or extrinsic regulations of underlying systems. It is unclear whether bimodality occurs in the absence of these regulations. The CTM has made great success in interpreting many experimental phenomena, but it cannot capture the bimodal distributions with two nonzero peaks that have been demonstrated in experiments. In particular, recent single-cell studies have shown non-exponential inactive periods and non-geometric translational bursting in gene expression. How to model these kinetics is challenging. We develop a stochastic gene model, namely the GTM, to model the silent transcription intervals by a general waiting-time distribution and translational bursting by an arbitrary distribution. By mapping the GTM into a queuing model, we derive the steady-state distribution of gene products that can be used for analyzing phenotypic switching. We find that non-exponential inactive times and translational bursting can lead to two nonzero bimodalities that cannot be captured in the CTM. These results indicate that both silent transcription intervals and translational bursting have important roles in controlling cell phenotypic variations in fluctuating environments.

## I. INTRODUCTION

Phenotypic switching, which is dominated by the stochastic fluctuation of gene expression, is an effective strategy for cells making decisions in changing environments.^1–4^ Single-cell experiments have provided evidence for bimodality of gene expression as a mechanism generating phenotypic diversity in a population of genetically identical cells.^5–7^ Each maximum of bimodality is usually associated with a subpopulation of a particular phenotype, which has been observed in stem cells,^8, 9^ and cancer cells.^10, 11^ Therefore, revealing the mechanism of phenotypic switching or bimodality is an important step in understanding fundamental cellular functions and variations.

Several viewpoints have been proposed to identify the mechanisms of bimodal gene expression. One common view is that bimodality results from deterministic bistability, i.e., two deterministic stable steady states of a system in the absence of stochasticity. Gene regulatory systems that produce this type of bimodality consist of a single positive feedback loop with cooperativity^12, 13^ [e.g., bacteriophage *λ*,^14^ Lactose operon of *Escherichia coli*^15^], or multiple feedback loops without cooperativity,^16^ and usually are accompanied by gene-state switching.^17^ The second viewpoint is noise-induced bimodality, i.e., the stochastic fluctuation can induce a bimodal response that cannot occur in the deterministic case.^5–7, 18, 19^ For example, using a synthetic system in budding yeast, To *et al*. found that positive feedback involving a promoter with multiple transcription factor (TF) binding sites can induce bimodality without cooperative binding of the TFs.^5^ The third viewpoint is noise filter-induced bimodality, which involves a nonlinear noise filter characterized by a Hill function that can transform the distribution of transcription rates from unimodal to bimodal in a cell population.^20^ The above three mechanisms of bimodality have similar features, that is, the underlying biochemical systems mostly involve intrinsic or extrinsic regulations, e.g., noise or feedback.^21–23^ A natural question is whether the bimodality of gene expression can occur in the absence of these regulations.

The most widely used model that reveals the molecular mechanisms of gene expression is the classical telegraph model (CTM),^24, 25^ where the promoter is assumed to switch between active (ON) and inactive (OFF) states, with only the former permitting transcription initiation. The CTM model elucidates the mechanisms of gene expression that do not involve intrinsic or extrinsic regulations, and has made great success in interpreting many experimental phenomena.^26–28^ Especially, it can explain the experimentally observed bimodal distributions of gene products that cannot be captured by the one-state model.^29–31^ Note that the distributions of gene products in the CTM only follow three types: unimodal located in origin, unimodal away from the origin, bimodal with one peak at origin while another peak away from the origin.^32^ However, some experimental studies have shown that the distributions of gene products may display two nonzero bimodalities,^33, 34^ e.g., hundreds of key immune genes are bimodally expressed with two nonzero maxima across cells.^7^ As we know, this type of bimodality cannot occur in the CTM, because the CTM could oversimplify the complex processes of gene expression such as gene-state switching and transcription or translation.

In fact, the process of stochastic gene-state switching may be very complex, e.g., transcription initiation involving the ordered assembly of pre-initiation complexes, recruitment of various polymerases, and chromatin remodeling, etc.^35, 36^ There have been studies showing that the complex process of stochastic gene-state switching is crucial for the change of cell phenotype switching.^2, 37–39^ Recent analysis of transcriptional dynamics in yeast and mammalian cells indicates that gene-state switching from an inactive state to the active state involving multiple slow biochemical reactions, which lead to non-exponentially distributed silent transcription intervals.^40–43^ For example, the heterogeneous inactive times give rise to the widely observed “noise” in human gene expression and explain the distribution of protein levels in human tissue.^44^

Given the complexity of silent transcription, how to model the process is challenging. One possible way is to consider multiple intermediate states, i.e., the promoter architecture involving multiple OFF or ON states.^45–49^ Although the increasing numbers of gene states can improve the fit between experimental data and theoretical models,^43, 44^ the difficulty in determining the numbers of promoter states as well as kinetics parameters would be detrimental to the inference of the data.^50–52^ Another alternative way is to use a non-Markovian modeling framework by introducing general dwell-time distributions for OFF or ON states.^53^ In this framework, the complex gene expression processes are often mapped into continuous-time random walk models,^54^ or queuing theory models.^55–60^ For example, Kumar *et al*. derived the analytical expressions of the steady-state moments for estimation of burst parameters by a single-state model.^58^ Shi *et al*. derived the analytical distribution of gene product in a two-state model with non-exponentially distributed inactive periods.^59^ Latter, Zhang *et al*. extended this model to double activation pathways (i.e., crosstalk), and derived the exact expressions for steady-state mRNA distribution and noise.^60^ The analysis of these models is mainly confined to steady-state moments and noise of protein counts, but the precise role of silent transcription times regulating the transition among the phenotypic states in a single cell is not understood.

The stochastic gene-state switching gives rise to transcriptional events occur in a bursty fashion, with the synthesis of mRNAs in short periods followed by longer silent intervals.^27, 40, 52, 61–64^ At the same time, single-cell experiments have provided evidence that the translation output of individual mRNA is highly variable and sporadic, resulting in the synthesis of protein in bursts, i.e., translational bursting.^65–68^ These bursting can lead to variations in gene expression levels and phenotypic in a population of genetically identical cells.^69^ In fact, translation is a complex biochemical process involving multiple factors and rate-limiting steps,^70^ implying that translational burst-size distribution would not be a geometric distribution.^58, 71^ For example, an Erlang distributed mRNA lifetime would lead to a negative binomial distributed burst size,^72^ whereas a deterministic constant mRNA lifetime would make the burst size obey a Poisson distribution.^73^ However, it is unclear how the translational burst-size distribution influences phenotypic switching and gene expression dynamics.

The above analysis shows the important roles of silent transcription intervals and translational bursting in regulating gene expression. Still, a challenging task is how to couple these two factors into a gene expression system. In this study, we develop a generalized telegraph model (GTM) of bursty gene expression, which considers general waiting-time distribution describing silent transcription intervals and arbitrary burst distribution characterizing translational bursting. With the queuing theory, we derive the binomial moment and gene product distribution that can be used for investigating phenotypic switching. We decompose the total protein noise into three parts: the low copy–number noise due to probabilistic individual birth and death events, the waiting-time noise resulting from stochastic gene-state switching between ON and OFF states, and the noise resulting from translational bursting. As a result, we show that the GTM can produce two nonzero bimodalities of protein distribution, implying that non-exponential inactive times and translational bursting have essential roles in regulating phenotypic switching. In addition, we find that both silent-intervals noise (i.e., OFF time noise) and burst-size noise can amplify gene expression noise. Finally, we explore the effects of silent transcription intervals and translational bursting on time-dependent gene expression dynamics, and find that silent-intervals noise and burst-size noise can induce diverse dynamic expressions. Our results not only provide an alternative mechanism for phenotypic switching driven by inactive periods and translational bursting, but also provide signatures useful for inferring complex promoter structure based on experimental data.

## II. MODEL AND METHODS

### A. Model description

We consider a general model of gene expression, called GTM, as outlined in Fig. 1. This model assumes that the gene promoter has one inactive (OFF) state and one active (ON) state, and the waiting-time distributions for the promoter dwelling at the OFF and ON state are denoted by *f*_off_(*t*) and *f*_on_(*t*), respectively. Protein is produced at the constant rate *r*_syn_ in an instantaneous bursting manner when the gene is activated.^29, 74^ Protein degradation rate is also a constant rate denoted by *r*_deg_. Based on these assumptions, we list all involved biochemical reactions as follows

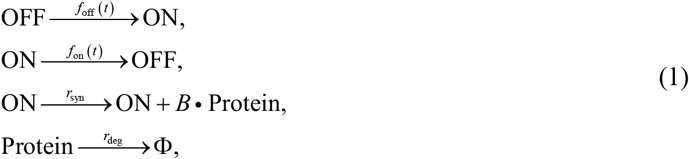

where *B* is the burst size, referring to the number of proteins created to be a burst distribution (see Fig. 1). Note that this model is an extension of previously studied models.^59, 60^ In addition, if *f*_off_(*t*) and *f*_on_(*t*) are exponentially distributed, then the gene model described by Eq. 1 reduces to a two-state model with translational bursting.^49^ Further, if burst size is the constant distribution with *B* ≡ 1, then Eq. 1 reduces to the CTM.^24, 25^ Here, we assume that the active-intervals (i.e., ON time) distribution in the GTM is an exponential distribution denoted by 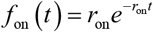 with *r*_on_ being promoter transition rate from ON state to OFF state, and the silent-intervals (i.e., OFF time) distribution *f*_off_(*t*) is a general distribution, which has been observed in recent experiments.^40–43^

**FIG. 1.**
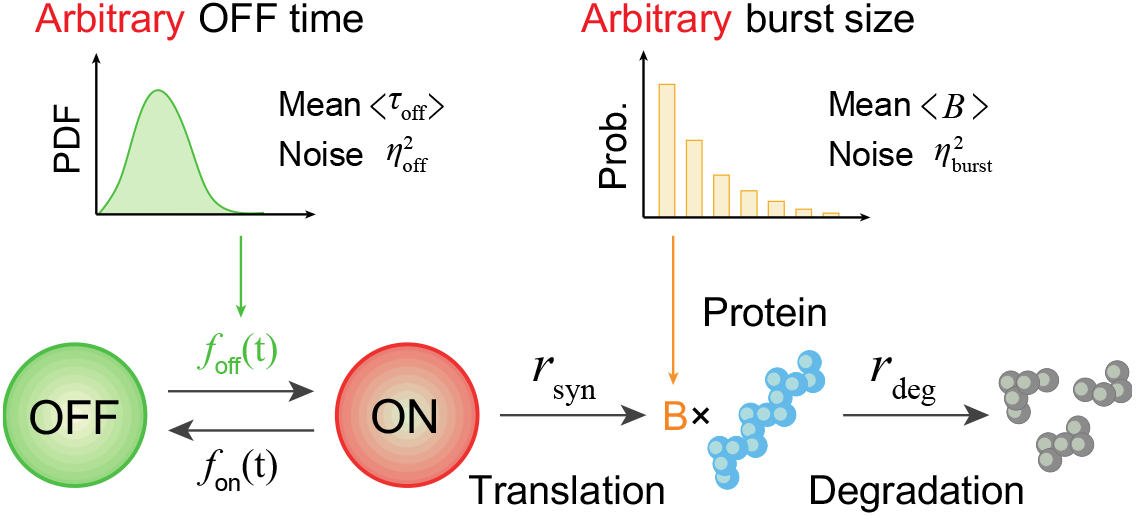
Schematic diagrams for the generalized telegraph model (GTM), where the OFF time follows an arbitrary distribution denoted by *f*_off_(*t*) with mean 〈*τ*_off_〉 and noise 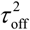, as well as the translational burst size follows an arbitrary distribution characterized by a random variable *B* with mean 〈*B*〉 and noise 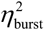. In addition, the ON time distribution is represented by *f*_on_(*t*), and the rates of synthesis and degradation of protein are constants denoted by *r*_syn_ and *r*_deg_, respectively.

### B. Mapping to queuing model

To explore the effects of general silent intervals and translating bursting on gene expression, we map the GTM to a queuing model: the production of proteins in bursts corresponds to the arrival of customers in batches, and the waiting-time distribution for protein decay corresponds to the service-time distribution for customers. Based on the mapping, the gene model represented by Eq. 1 can be described as a *GI^X^*/*M*/∞ queuing model.^55, 58, 75^ Here, *G* refers to a general waiting-time distribution between two successive burst events of protein, which is a very critical quantity for analysis of queuing systems. For convenience, we call initiation-time distribution, and denote it by *f*_ini_(*t*) in our study. *I^X^* denotes protein producing in batches of independently distributed random size *X* (i.e., burst size *B*), and *M* represents Markovian (i.e., exponential) degradation-time distribution for protein. In addition, ‘∞‘ means that protein degradation is independent and is not affected by protein creation. For this queuing model, as far as we know, no exact results such as the steady-state distribution have been reported. Applying the mapping, we will derive the steady-state protein distribution of the GTM for analyzing phenotypic switching.

### C. Initiation-time distribution and renewal function

According to the above analysis of queuing model, the key step for computing the exact protein distribution shown in Eq. 1 is to obtain the analytical expression for the initiation-time distribution *f*_ini_(*t*). From Fig. 1, we note that the promoter in the ON state can either produce protein bursting or switch back to OFF state. More specifically, the gene in the ON state can produce protein bursting in a single step, or make multiple trips to the OFF state before producing protein bursting. As discussed in a previous study, a calculation based on this set-up can lead to the following explicit expression for the initiation-time distribution in the Laplace domain^58^

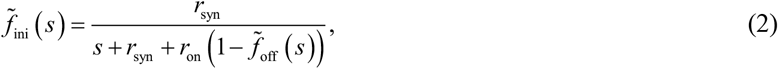

where 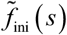 and 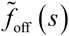 are the Laplace transforms of *f*_ini_(*t*) and *f*_off_(*t*), respectively. In particular, if we define the survival function of the OFF state 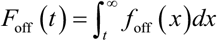 with its Laplace transform denoted by 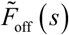, and then it is easy to find 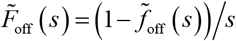. Substituting this expression into Eq. 2, we further obtain 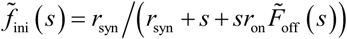. Finally, the initiation-time distribution *f*_ini_(*t*) can be obtained by calculating the inverse Laplace transform of 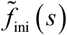, which is given by 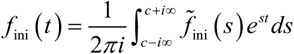.

Next, we calculate the analytical expression of the renewal function. The renewal function is defined as the mean number of renewal events. It is a key quantity in the field of queueing theory since most of the analytical results, such as the binominal moments and distribution, can be obtained based on the renewal function. The renew function *R* (*t*) can be given by the following renewal equation^56^

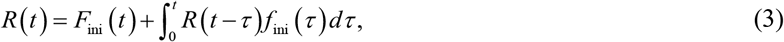

where *F*_ini_(*t*) is the cumulative distribution function for the initiation-time distribution, i.e., 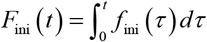. Applying the Laplace transform to Eq. 3, we have 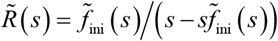, where 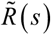 is the Laplace transform of the renewal function. Substituting the expression of 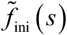 into 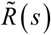, we further obtain

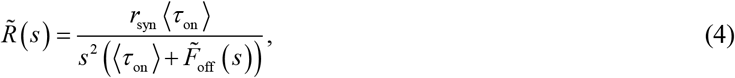

where 〈*τ*_on_〉 = 1/*r*_on_ is the mean ON time. Finally, the renewal function can be obtained by computing the inverse Laplace transform of 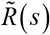, which is given by 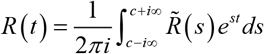.

### D. Binomial moments and probability distribution

To compute the statistic moments and distribution of gene product amounts, we apply binomial moment method.^47, 77^ The binomial moment is a key quantity for computing important statistical indices such as mean, noise, skewness, and kurtosis.^49, 60^ Let 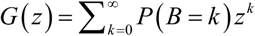 denote the generating function of the protein burst distribution. The *i*th binomial moment of the burst size *B*, denoted by 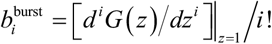 with 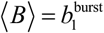 representing the mean burst size. Then the time-dependent binomial moment for our gene model can be given by^75^

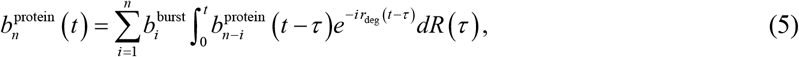

where 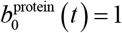 due to the probability conservation. Eq. 5 shows that all the time-dependent moments can be derived if the renewal function is analytically given. For example, the time-dependent protein average 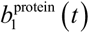 and noise 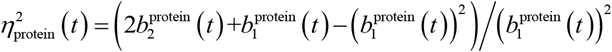 can be obtained based on the renewal function.

In addition, if we denote by the mean OFF time 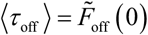, and the mean translational rate 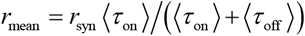, we further give the steady-state binomial moment

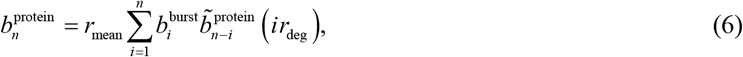

where the Laplace transform is given by the following relation

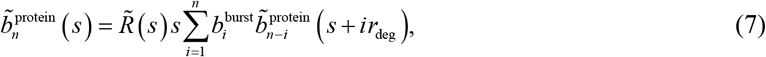

with 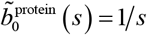 and 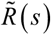 being given in Eq. 4. These results yield that all steady-state moments, including each order raw moment and center moment, can be obtained according to the Laplace transform of renewal function and the iterative relations in Eq. 6 and Eq. 7.

For clarity, we give analytical expressions for the first four steady-state binomial moments based on Eq. 6 and Eq. 7. The first-order binomial moment, i.e., the protein average, is given by

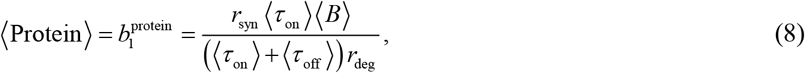

which implies that the mean protein level depends on the mean time of promoter dwelling in two states and average translational burst size. The second-order binomial moment takes the form

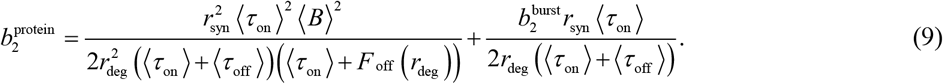

Thus, the protein noise at steady state can be quantified by

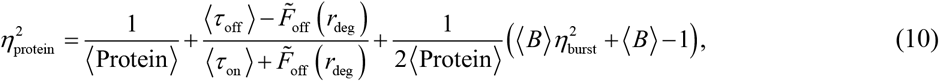

where 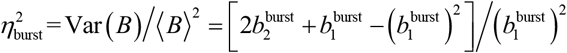 is the burst-size noise. Eq. 10 highlights different contributions to the protein noise: the low copy-number noise due to probabilistic individual birth and death events (the first term), the waiting-time noise resulting from stochastic promoter switching between two states (the second term), and translational bursting noise which is the function of mean burst size and burst-size noise (the third term).

The third-order binomial moment is given by

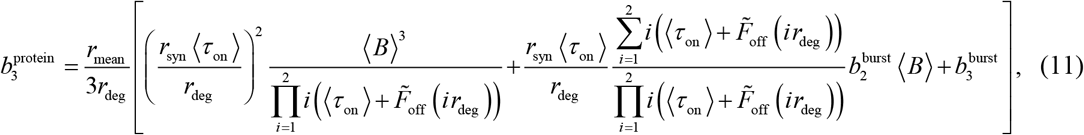

and the fourth binomial moment by

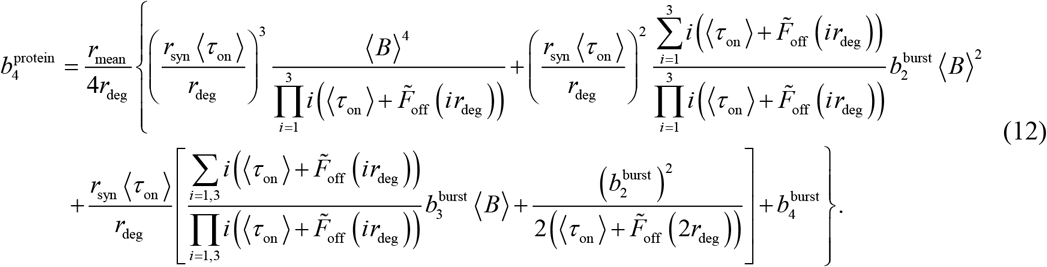

In addition, using the following relation between binomial moments and central moments

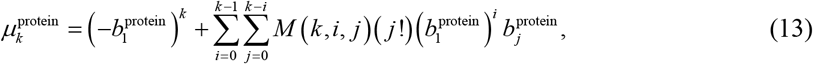

where 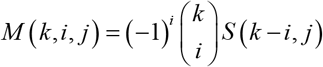 with 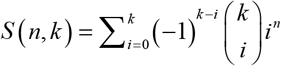 being the Stirling number of the second kind, we can further compute the skewness and kurtosis, which are denoted by 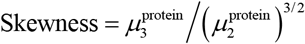 and 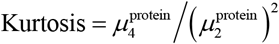, respectively.

In particular, the binomial moments can be used in the reconstruction of underlying protein distribution according to

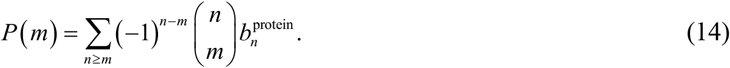

## III. EXACT RESULTS FOR SPECIAL CASES

In this section, we will give the exact steady-state distribution of protein abundance for representative silent-intervals distribution (e.g., Erlang distribution) and translational burst distribution (e.g., geometric distribution).

### A. Analytical results for initiation-time distribution and renew function

Here, we consider that the waiting-time distribution for silent transcription intervals is the Erlang distribution with 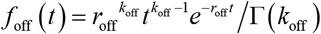, where *r*_off_ is the scale parameter and *k*_off_ is the shape parameter often used for characterizing the number of reaction steps.^54, 60^ Note that *k*_off_ = 1 corresponds to the case of exponential waiting time, whereas *k*_off_ > 1 corresponds to the case of non-exponential waiting time. In addition, the mean silent time is 〈*τ*_off_〉 = *k*_off_/*r*_off_. To derive the analytical expression for initiation-time distribution, we first compute the Laplace transform of the survival function *F*_off_(*t*). It is easy to get 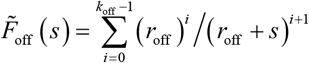. Substituting this expression into expression 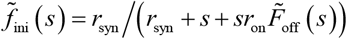, we further obtain the Laplace transform of initiation-time distribution *f*_in_(*t*), that is

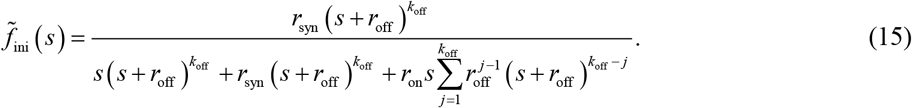

Note that the Laplace transform *f*_ini_ (*s*) in Eq. 15 can be given by the following rational function

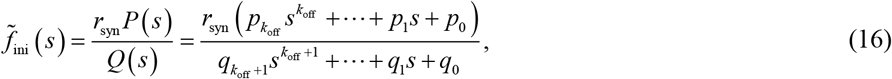

where *p_i_*(*i* = 1,2, ⋯, *k*_off_), and *q_j_*(*j* = 1,2, ⋯, *k*_off_ +1) are constant coefficients determined by Eq. 15. Assume that *Q*(*s*) has *l* real roots and *m* pairs of complex roots (i.e., *l* + 2*m* = *k*_off_ +1). By using the partial fraction expansion, 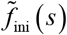 can be decomposed into the summation of real part and complex part

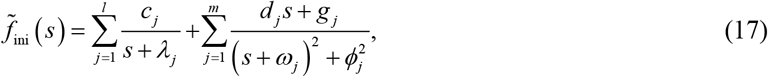

where coefficients *c_j_,d_j_,g_j_,λ_j_,ω_j_* and *ϕ_j_* are obtained from the partial fraction expansion. Using the inverse Laplace transform to Eq. 17, we obtain the following initiation-time distribution

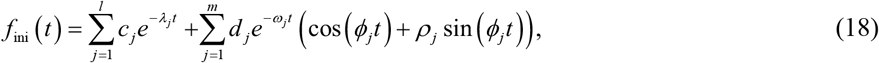

where 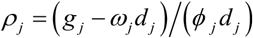. Eq. 18 implies that the analytical expression of initiation-time distribution can be obtained if its Laplace transform can be written the partial fractions.

Next, we derive the analytical results for the renewal function *R*(*t*). Substituting the expression 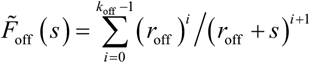 into Eq. 4, we obtain the Laplace transform of the renewal function *R*(*t*) as follows

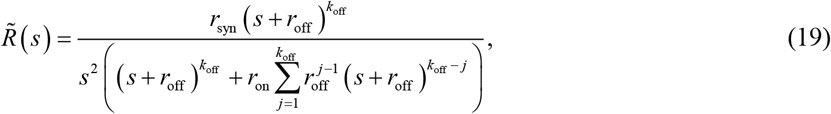

Note that the Laplace transform 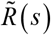 in Eq. 19 can be given by the following rational function

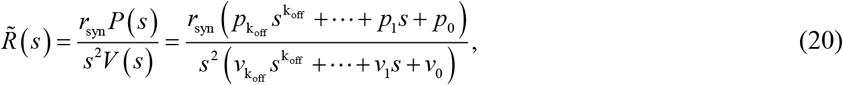

where *v_i_*(*i* = 1,2, ⋯, *k*_off_) is constant-coefficient determined by Eq. 19, and *p_i_*(*i* = 1,2, ⋯, *k*_off_) has the same expression as in Eq. 16. Assuming *V*(*s*) has *l*_1_ real roots and *m*_1_ pairs of complex roots (i.e., *l*_1_ + 2*m*_1_ = *k*_off_). By using the partial fraction expansion, 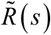 can be decomposed into the summation of real part and complex part

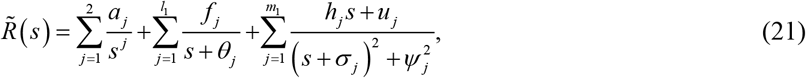

where coefficients *a_j_*, *f_j_*, *h_j_*, *u_j_*, *θ_j_*, *σ_j_*, and *ψ_j_* are obtained from the partial fraction expansion. Using the inverse Laplace transform to Eq. 21, we obtain the renewal function

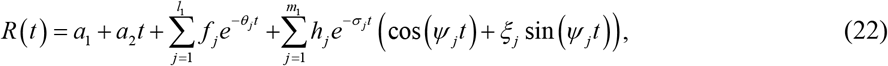

where 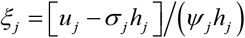. Eq. 22 implies that the analytical renewal function can be obtained if its Laplace transform can be written the partial fractions.

Substituting Eq. 22 into Eq. 5, we obtain the time-dependent first-order binomial moment, i.e., the time-dependent protein average 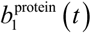, as follows

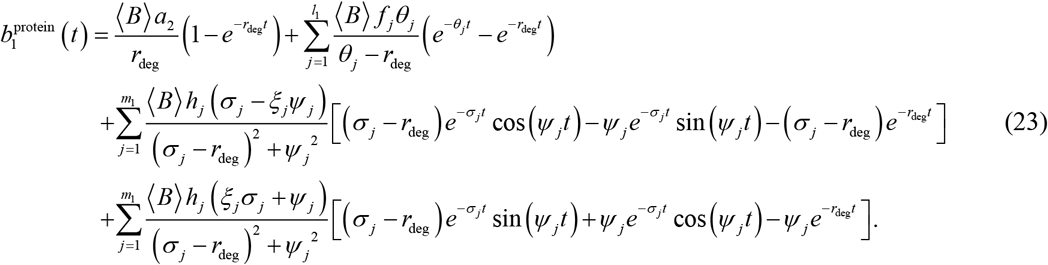

In addition, we can also give the analytical expressions for a time-dependent second-order binomial moment 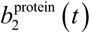 and time-dependent noise 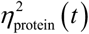, but we omit them due to the complexity of their expressions.

### B. Analytical results for steady-state binomial moments and probability distributions

Here, we consider two representative protein burst distributions: (1) Constant distribution with *P*(*B* = 1) = 1, *P*(*B* = *k*) = 0, *k* = 1,2,3, ⋯; (2) Geometric distribution with 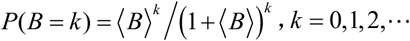. For each burst distribution, we derive the analytical expression of steady-state binomial moments and probability distributions.

#### Case 1: Constant distribution for burst size

For constant burst-size distribution, we have the binomial moment 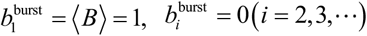. Substituting the expression of 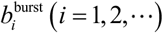 into Eqs. 6 and 7, we obtain the following expression

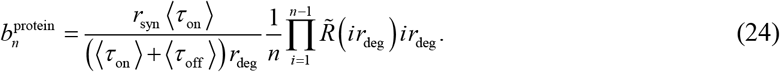

Combining Eq. 24 with the expression of *R*(*s*) in Eq. 20, we further obtain the analytical expression for the steady-state binomial moments

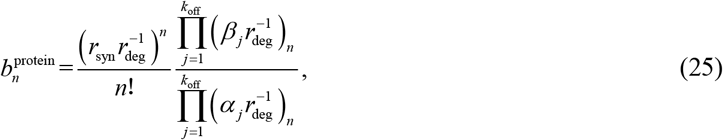

where −*β_j_* and −*α_j_*(*j* = 1,2, ⋯, *k*_off_) are the roots of Equations 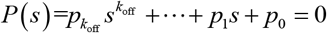 and 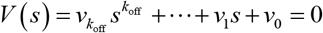, respectively. In addition, symbol (*υ*)_*n*_ =*υ*(*υ* – 1)⋯(*υ*–*n* +1), *n* = 1,2, ⋯. Substituting Eq. 25 into Eq. 14, we obtain the steady-state protein distribution

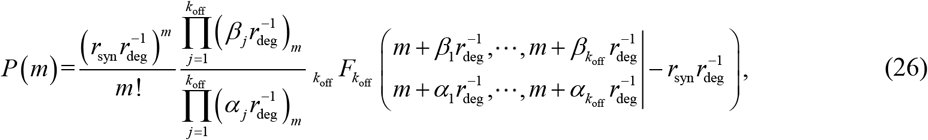

where _*k*_off__*F*_*k*_off__ denotes a generalized hypergeometric function. In particular, for the exponential waiting time, i.e., 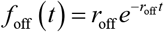, we have the analytical expression of the protein distribution as follows

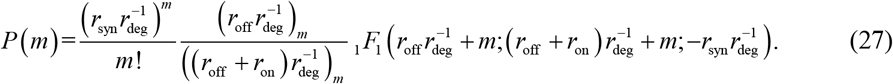

This distribution has been derived in previous works.^46, 47^

#### Case 2: geometric distribution for burst size

For geometric burst-size distribution, we have the binomial moment 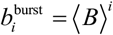. Substituting the expression 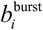 into Eqs. 6 and 7, we obtain the following expression

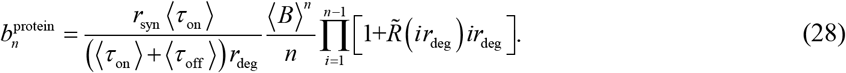

Combining Eq. 28 with the expression of *R*(*s*) in Eq. 20, we further obtain the analytical expression for the steady-state binomial moments

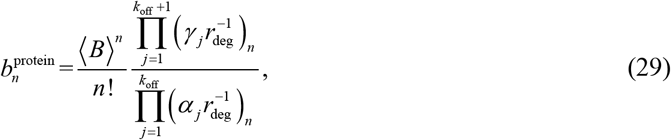

where −*γ_j_* (*j* = 1,2, ⋯, *k*_off_ +1) are the roots of 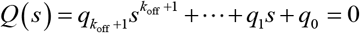, and *α_j_* (*j* = 1,2, ⋯, *k*_off_) have the same expressions as in Eq. 25. Substituting Eq. 29 into Eq. 14, we further obtain the steady-state protein distribution

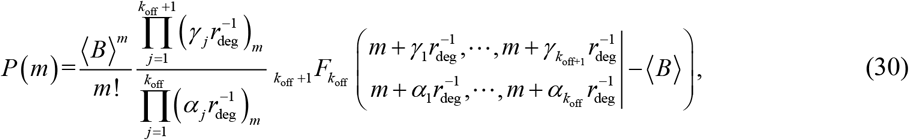

where _*k*_off_ +1_*F*_*k*_off__ denotes a generalized hypergeometric function.

In particular, for the exponentially distributed OFF waiting time, i.e., 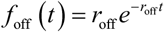, we can obtain the analytical expression of steady-state protein distribution as follows

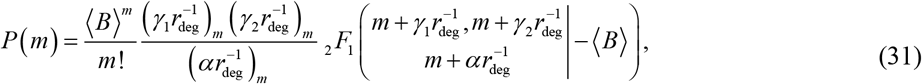

where 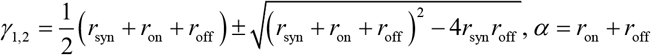. This distribution has been derived in previous work.^49^

## IV. NUMERICAL RESULTS

In this section, we numerically demonstrate the influences of both silent transcription intervals and burst stochasticity on gene expression. For this, we introduce two indices: switching correlation time 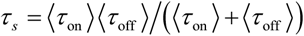 and promoter mean occupancy 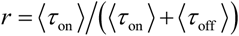, which can be used for fluctuation analysis in gene expression across a range of parameters.^64^

### A. Non-exponential inactive times can lead to two nonzero bimodalities of protein distribution

First, we explore the effects of silent transcription intervals on protein distribution. To map how the shape of protein distribution varies with switching correlation time *τ_s_* and promoter mean occupancy *r*, we define the bimodality coefficient (BC):^78^ BC = 1/(Kurtosis – Skewness^2^), where 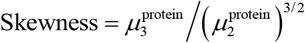 and 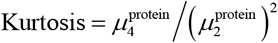 with the central moment 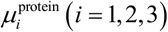 obtained according to Eq. 13. The threshold of BC greater than 5/9 may indicate a bimodal or multimodal distribution. Calculating BC over a broad range of kinetics rates, we find that the protein distribution is shaped by *τ_s_* and *r*, referring to Fig. 2.

**FIG. 2.**
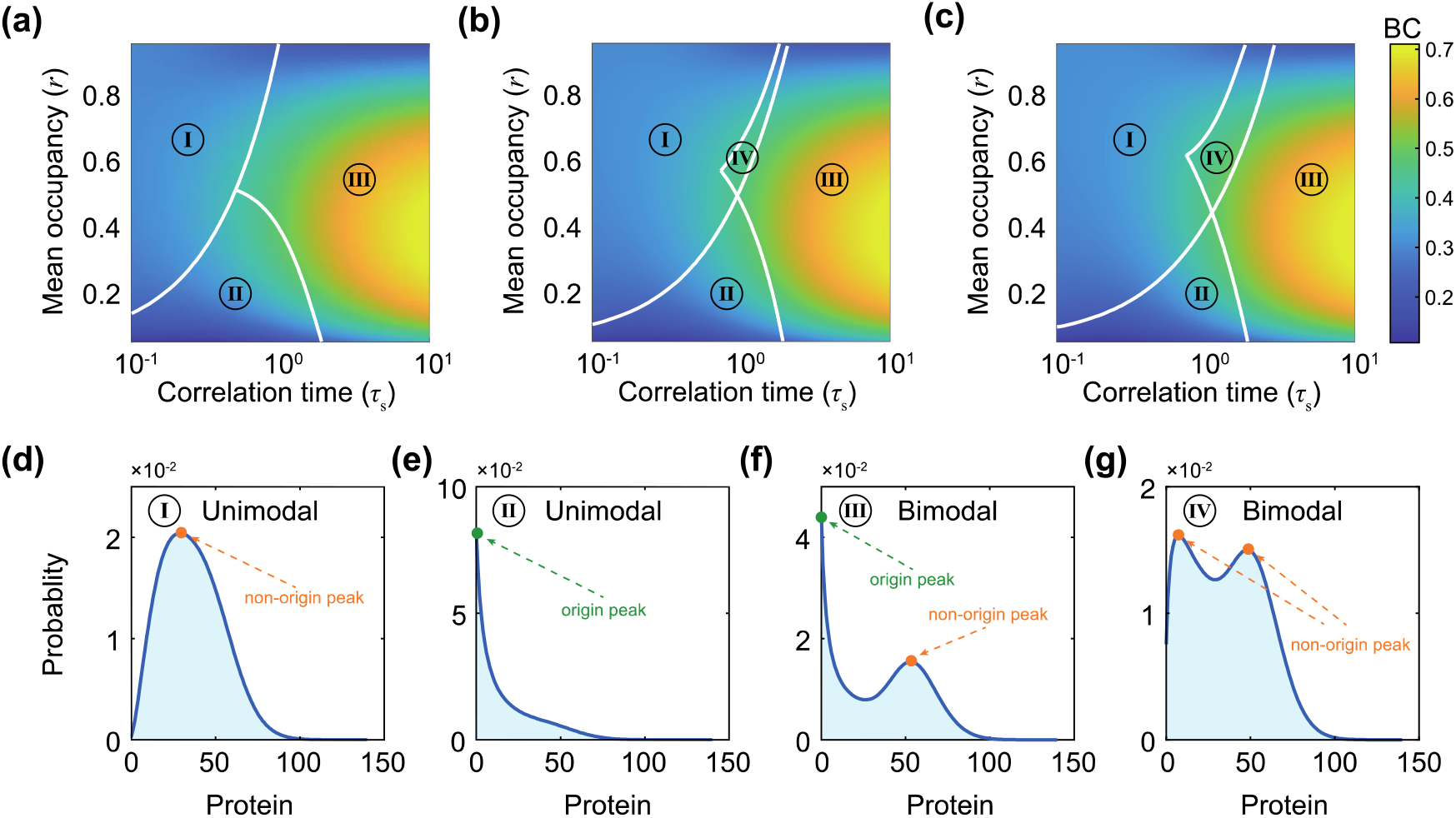
Effects of silent transcription intervals on protein distribution in the *τ_s_* – *r* plane. (a-c): Heatmap, respectively, the bimodality coefficient (BC) as a function of switching correlation time *τ_s_* and promoter mean occupancy *r* to classify protein distribution for three different silent-intervals noise of Erlang distribution: 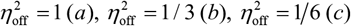. We denoted by I, II, I, and IV the corresponding parameter regions to the unimodal located in origin, unimodal away from the origin, bimodal with one peak at origin while another peak away from the origin, and bimodal with two peaks away from the origin. (d-g): Shown are representative protein distributions corresponding to regions I, II, I, and IV, respectively. Parameters are set as: *r*_syn_ = 30, *r*_deg_ = 1, as well as *r*_off_ and *r*_on_ are calculated by *r*_off_ = *k*_off_*r*/*τ_s_*, *r*_on_ = (1 – *r*)/*τ_s_*, respectively. In addition, the burst-size distribution is chosen as geometric distribution with fixed burst-size noise 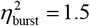.

Fig. 2 shows the influences of silent transcription intervals on the classification of protein distribution with the modulation of switching correlation time and promoter mean occupancy. Here, we consider the OFF waiting-time distribution as the Erlang distribution: 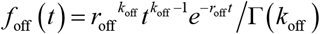. Accordingly, the mean silent time can be given by 〈*τ*_off_〉 = *k*_off_/*r*_off_, and silent-intervals noise can be denoted as 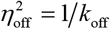. In addition, the burst-size distribution is assumed to be geometric distribution with fixed burst-size noise. We choose the silent-intervals noise 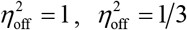 and 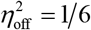, referring to Fig. 2(a), 2(b), and 2(c), respectively. That is, Fig. 2(a) corresponds to the case of exponentially distributed silent transcription intervals, whereas Figs. 2(b) and 2(c) correspond to the case of non-exponentially distributed silent transcription intervals. From Fig. 2(a), we observe that there are only three modes for protein distribution: unimodal located in origin (denoted by I), unimodal away from the origin (denoted by II), and bimodal with one peak at origin while another peak away from the origin (denoted by III). However, we find that there are bimodal with two nonzero peaks (denoted by IV) in the case of non-exponentially distributed silent transcription intervals, referring to Figs. 2(b) and 2(c). In particular, the bimodal region of type IV was further expanded as the further enhancement of the non-exponential behavior (comparing Fig. 2(b) with Fig. 2(c)). Finally, four representative distributions corresponding to regions I to IV are shown in Fig. 2(d)–2(g), respectively. These results indicate that non-exponential inactive times can lead to two nonzero bimodalities of the proteins’ steady-state distribution.

Previous studies showed that bimodal gene expression is a cause of phenotypic diversity in genetically identical cell populations.^5–7^ On the other hand, recent experiments have shown that the inactive periods for many genes have a non-exponential distribution in mammalian cells.^40–43^ Our results suggest that the non-exponential behaviors can significantly enhance the bimodality such as expanding the region of two nonzero peaks. Thus, we speculate that the non-exponentially distributed inactive period would be a cause of achieving phenotypic diversity, which is critical for cell survival in a fluctuating environment.^69^

### B. Larger burst size can induce bimodal protein distribution

In this subsection, we analyze the influences of burst size on gene product distribution. Here, we consider geometrically distributed burst size with probability distribution 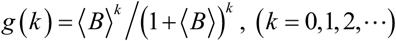 and burst-size noise 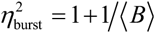. In addition, the OFF-time distribution is assumed to be the Erlang distribution. Numerical results are shown in Fig. 3, where we change the mean burst size and fix the switching correlation time and promoter mean occupancy. Fig. 3(a) shows how the number of the most probable protein molecules depends on mean burst size, where pink denotes the bimodal region. We observe that the shapes of the protein distribution change from unimodal to bimodal, and the distance between the two non-origin peaks of bimodal distribution become larger as the mean burst size increases, implying that burst size can induce the bimodal distribution of gene product. Fig. 3(b)-(d) show three representative protein distributions corresponding to the letter b, c, and d in Fig. 3(a), respectively, where solid lines are theoretical predictions, whereas circles are stochastic simulations.^79^ These results imply that translational bursting plays a significant role in generating phenotypic diversity, which has been verified in recent single-cell and single-molecule experiments.^33^

**FIG. 3.**
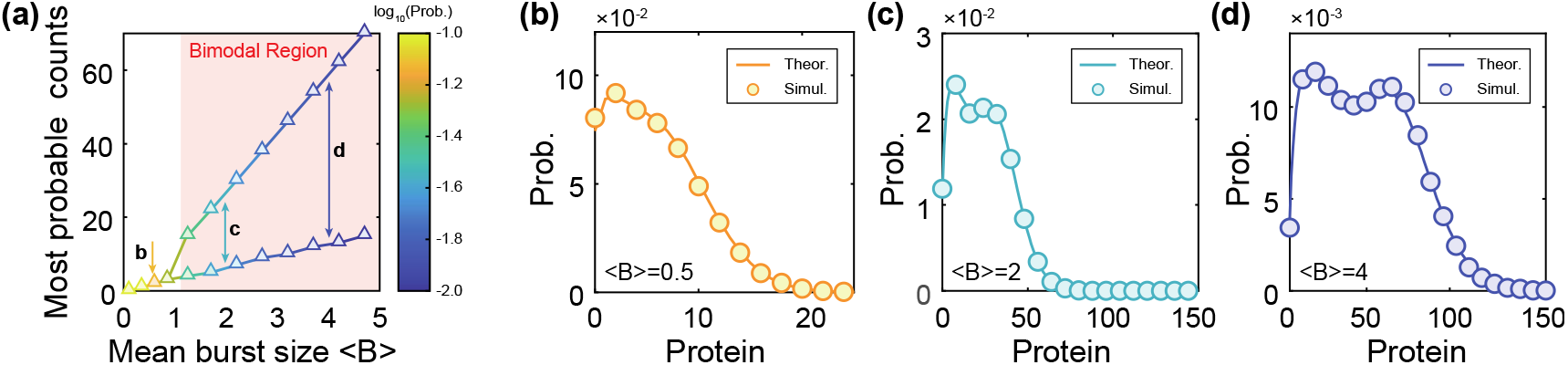
Influence of burst size on protein distribution. (a) Dependence of the most protein counts on mean bust size, where the pink region denotes the presence of bimodality, and the letter b, c, and d corresponds to mean burst size 〈*B*〉 = 0.5, 2,4, respectively. (b) The case where the protein distribution is unimodal with one nonzero mode when mean burst size 〈*B*〉 = 0.5. (c) and (d) The cases where the protein distribution is bimodal with two nonzero modes when the mean burst size 〈*B*〉 = 2 and 〈*B*〉 = 4, respectively. Here, the burst size is assumed to be geometric distribution, and OFF time is considered as Erlang distribution with fixed silent-intervals noise 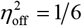. The parameters are set as: *r*_syn_ = 20, *r*_deg_ = 1, *k*_off_ = 6, *r*_off_ = 3.6, *r*_on_ = 0.4.

### C. Both silent-intervals noise and burst-size noise can amplify gene expression noise

First, we examine the effects of silent transcription intervals on gene expression dynamics and consider the waiting time for the inactive periods as Erlang distribution, e.g., 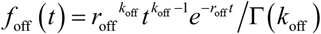 with mean silent time 〈*τ*_off_〉 = *k*_off_/*r*_off_, and silent-intervals noise 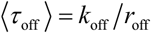. In addition, we assume the burst-size distribution as a geometric distribution with fixed burst-size noise 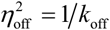. Numerical results are demonstrated in Fig. 4(a), where we vary the promoter mean occupancy *r* while keeping the switching correlation time *τ_s_* constant. From Fig. 4(a), we observe that larger silent-intervals noise can produce more protein noise, implying that the dynamics of inactive duration can be propagated into protein and encode the stochastic dynamics of the protein.

**FIG. 4.**
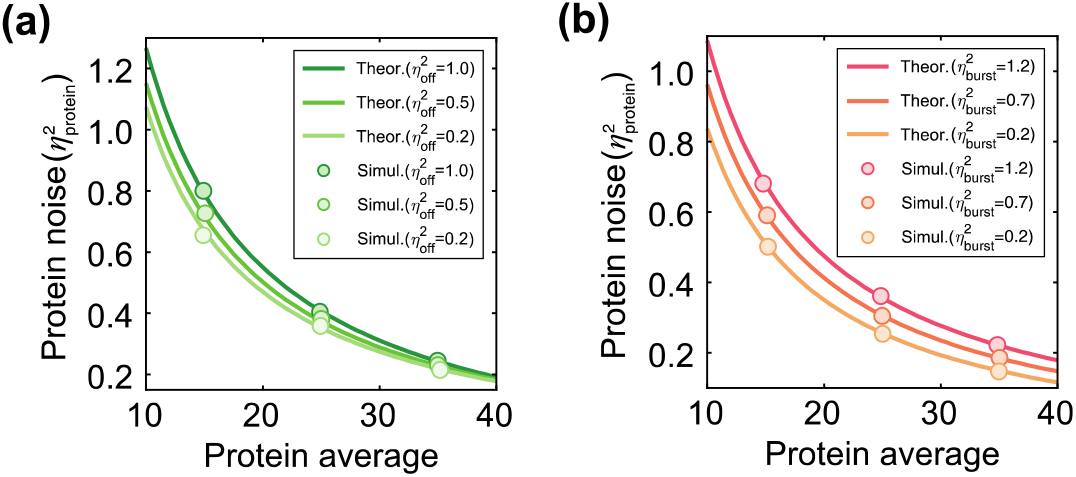
Effects of silent-intervals noise and burst-size noise on protein fluctuations for varying the promoter mean occupancy *r* while fixing the switching correlation time *τ_s_*. (a) Noise versus average in protein abundance for three different silent-intervals noise 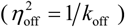 from Erlang distribution. The parameters are set as: *r*_syn_ = 10, *r*_deg_ = 1, *τ_s_* = 0.2, as well as *r*_off_ and *r*_on_ are calculated by *r*_off_ = *k*_off_*r*/*τ_s_*, *r*_on_ = (1 – *r*)/*τ_s_*, respectively. In addition, the burst-size distribution is chosen as geometric distribution with fixed burst-size noise 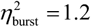. (b) Noise versus average in protein abundance for three different burst-size noise from geometric distribution 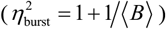, negative binomial distribution 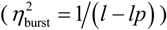, and Poisson distribution 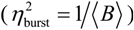, respectively. The parameters are set as: 〈*B*〉 = 5, *l* = 2, *p* = 2/7, and other parameters are the same as in case (a). In addition, the OFF waiting-time distribution is chosen as the Erlang distribution with fixed silent-intervals noise 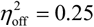. In (a) and (b), curves are theoretical predictions from Eq. 10, whereas circles are stochastic simulations.^79^

Then, we explore the effects of bursting kinetics on gene expression, and consider three different burst-size distributions: (i) Geometric distribution with probability distribution 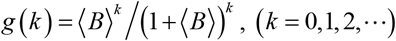 and burst-size noise 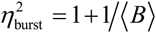. (ii) Negative binomial distribution with probability distribution 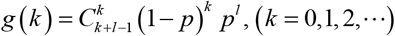 and burst-size noise 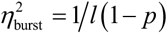. (iii) Poisson distribution with probability distribution 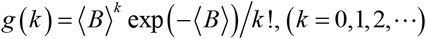 and burst-size noise 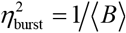. In addition, the OFF waiting-time distribution is considered to be Erlang distribution with fixed silent-intervals noise 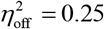. Numerical results are demonstrated in Fig. 4(b), where we modulate the promoter mean occupancy *r* by fixed switching correlation time *τ_s_*. From Fig. 4(b), we observe that larger burst-size noise can lead to more protein noise, implying that the translational burst kinetics can be propagated into protein and encode the dynamics of stochastic gene expression.

To summarize, Fig. 4 shows that both silent-intervals noise and burst-size noise can amplify gene expression noise, implying that the inactive times and translational bursting have important influences on fluctuations in gene expression. In addition, we find that these results in Fig. 4 support the canonical mean-noise inverse correlation across wide parameters.^80, 81^

### D. Silent-intervals noise and burst-size noise can lead to diverse dynamic expressions

The major difference between our GTM and the CTM is that the promoter switching from OFF state to ON state would involve intermediate reaction steps leading to non-exponential distributed silent transcription intervals. In addition, the GTM considers translational bursting, verified in experiments.^65–68^ Here, we explore the effects of silent-intervals distribution and burst-size distribution on the time-dependent average and gene expression noise. For the former, we consider silent intervals as Erlang distribution when burst size is assumed to be geometric distribution with fixed burst-size noise. For the latter, we consider three different burst-size distributions, i.e., geometric distribution, negative binomial distribution, and Poisson distribution, when OFF waiting time is taken as Erlang distribution with fixed silent-intervals noise. Numerical results are shown in Fig. 5, where Fig. 5(a) (Fig. 5(b)) plots the protein average (protein noise) versus time for three different silent-intervals noise, whereas Fig. 5(c) (Fig. 5(d)) plots the protein average (protein noise) versus time for three different burst-size noise.

**FIG. 5.**
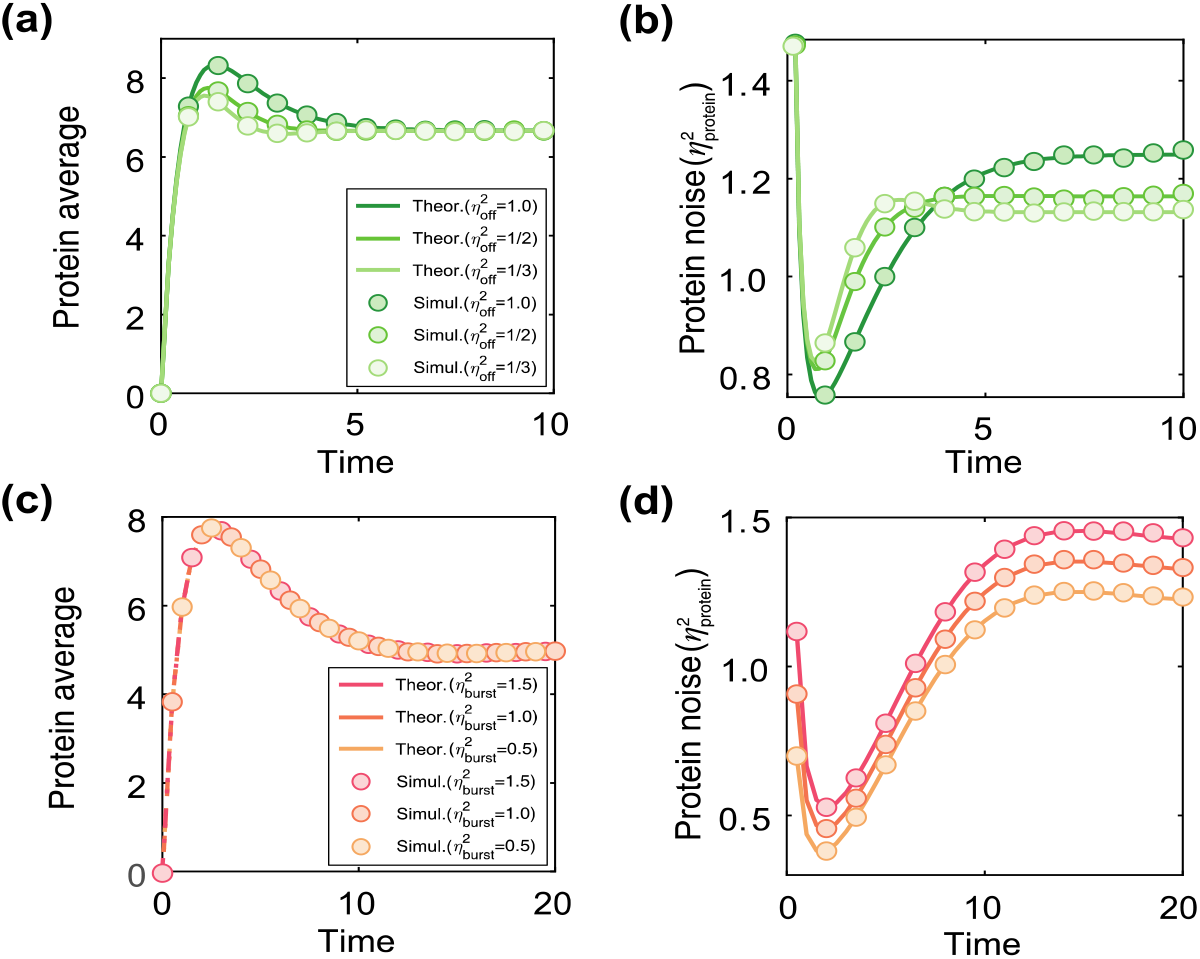
Influences of silent-intervals noise and burst-size noise on time-dependent average and noise of protein. (a) and (b) Shown are the protein average and noise versus time, respectively, for three different silent-intervals noise 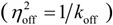 from Erlang distribution. The parameters are set as: *r*_syn_ = 10, *r*_deg_ = 1, *τ_s_* = 2/3, *r* = 1/3, as well as *r*_off_ and *r*_on_ are calculated by *r*_off_ = *k*_off_*r*/*τ_s_*, *r*_on_ = (1 – *r*)/*τ_s_*, respectively. In addition, the burst-size distribution is chosen as geometric distribution with fixed burst-size noise 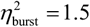. (c) and (d) Shown are the protein average and noise versus time, respectively, for three different burst-size noise from geometric distribution 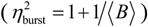, negative binomial distribution 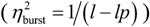, and Poisson distribution 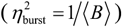, respectively. Here, the OFF waiting-time distribution is chosen as Erlang distribution with fixed silent-intervals noise 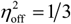. The parameters are set as: *r*_syn_ = 5, *r*_deg_ = 1, *k*_off_ = 3, *r*_off_ = 0.3, *r*_on_ = 0.1, 〈*B*〉 = 2, *l* = 2, *p* = 1/2. In all cases, curves are theoretical predictions, whereas circles are stochastic simulations.^79^

From Fig. 5(a), we observe that the time-dependent protein average has obvious differences for different silent-intervals noise, although long time (i.e., steady-state) average behavior is identical. However, we found that the time-dependent protein average is identical for three different burst-size noise, referring to Fig. 5(c). Then we analyze the influences of silent-intervals noise and burst-size noise on time-dependent protein noise. From Fig. 5(b), we find the time-dependent protein noise modulated by inactive periods shows complex characteristics: the lowest (largest) silent-intervals noise has the largest (lowest) protein noise for a short time, but there is an opposite relationship as the time further increase, i.e., larger silent-intervals noise can lead to larger protein noise, which is consistent with the steady-state result shown in Fig. 4. From Fig. 5(d), we find that time-dependent protein noise modulated by burst-size noise show the same characteristic as the steady-state behavior, i.e., larger burst-size noise can lead to larger protein noise. These results show that silent transcription intervals and translational bursting have important influences on dynamic gene expression.

## V. CONCLUSION AND DISCUSSION

Bimodal gene expression is considered a cause of phenotypic diversity in genetically identical cell populations,5–7 but identifying the bimodality mechanisms of underlying systems are continuing challenge. Previous studies on bimodality mostly involve the intrinsic or extrinsic regulations, e.g., noise and feedback, of the underlying systems.^12–23^ Bimodality characterized by mixed distributions also has been observed in these gene models involving promoter leakage,^82^ and multiple exits of transcription.^47^ However, it is unclear that bimodality can occur without these conditions. The CTM has made great success in interpreting many experimental phenomena,^26–28^ especially it can explain the observed bimodality that cannot be captured by the one-state model.^29–31^ However, it cannot capture the bimodal distribution with two nonzero maxima that has been observed in recent single-cell experiments.^7, 33, 34^ On the other hand, several experimental studies have shown that the inactive periods of a promoter have non-exponentially distributed waiting times.^40–43^ At the same time, studies have demonstrated that the complex translational process can lead to non-geometric burst distributions.^71–73^ Thus, it is necessary to develop effective methods to explore the effects of both inactive periods and translational bursting on bimodal gene expression.

In this study, we have developed the GTM for analyzing the bimodality of gene expression, which incorporates the two key features mentioned above: general silent transcription intervals and arbitrary translational bursting kinetics. By mapping the GTM to the system studied in queuing theory, we derived analytical expressions for the steady-state distributions and any order moment of protein counts. We decomposed the total protein noise into three parts: low-copy noise, waiting-time noise, and translational noise. We find that, for Erlang distributed silent transcription intervals, the exact steady-state distributions of protein can be denoted by two types of generalized confluent hypergeometric functions: one is _*k*_off__ *F*_*k*_off__ type if protein burst distribution is constant, and the other is _*k*_off_ +1_*F*_*k*_off__ type if protein burst distribution is geometric. We should point out that many previously studied results are the particular cases of our analytical results obtained here, referring to references.^46, 47, 49, 59, 60^

By introducing two indices named switching correlation time and promoter mean occupancy,^64^ we numerically demonstrated the influences of both silent transcription intervals and protein burst kinetics on bimodal gene expression. We have explored robust regions of unimodality and bimodality for different OFF waiting times, and found that non-exponential inactive times can lead to two nonzero bimodalities of protein distribution. At the same time, we investigate the effects of translational bursting on distribution modes of the gene product and found that a larger burst size can induce bimodal distribution. These results indicate that our GTM can capture a type of bimodality with two nonzero peaks, which cannot occur in the CTM.^32^ Our results complement and enrich the relative conclusions on bimodal gene expression, which are useful for analyzing phenotypic diversity in a complex cellular process.

In addition, we found that both silent-interval noise and translational burst-size noise can amplify protein noise, implying that the kinetics of silent transcription and translational bursting can be propagated into protein and encode the stochastic dynamics of the protein. Finally, we explored the effects of silent transcription intervals and translational bursting on time-dependent behaviors for both average and noise of protein, and found that the two kinds of noise can lead to diverse dynamic expressions. For example, the silent-intervals noise can induce different time-dependent mean behavior even under the same steady-state mean behavior, implying that caution must be kept when one infers a gene system based on steady-state behavior.

Note that multistate models in previous studies are introduced for characterizing complex promoter structure.^45–49^ However, the difficulty in determining the numbers of promoter states and kinetics parameters would be detrimental for interpreting experimental phenomena. Thus, combining relative experimental results with our theoretical approaches can be used to investigate the dynamics of gene expression in complex cellular processes. Note that we omitted the process of mRNA production under the assumption that mRNA lifetime is much shorter than protein lifetime. It is worthy of further study on the combined effect of gene expression when both transcriptional busting and translational bursting are considered simultaneously. In addition, some intermediate processes of gene expression such as cell division,^83^ alternative splicing,^84^ and elongation,^85^ are neglected in our study. How details of these factors affect phenotypic switching is worth further investigation.

## ACKNOWLEDGMENTS

This work was supported by the National Key R&D Program of China [grant number 2021YFA1302500]; the Natural Science Foundation of P. R. China [grant numbers 12171494, 11601094, 11931019, 11775314]; the Key-Area Research and Development Program of Guangzhou, P. R. China [grant numbers 2019B110233002, 202007030004]; and the Guangdong Basic and Applied Basic Research Foundation [grant number 2022A1515011540].

